# Multi-Method Characterisation of the Human Circulating Microbiome

**DOI:** 10.1101/359760

**Authors:** E. Whittle, M.O. Leonard, R. Harrison, T.W. Gant, D.P Tonge

## Abstract

The term microbiome describes the genetic material encoding the various microbial populations that inhabit our body. Whilst colonisation of various body niches (e.g. the gut) by dynamic communities of microorganisms is now universally accepted, the existence of microbial populations in other “classically sterile” locations, including the blood, is a relatively new concept. The presence of bacteria-specific DNA in the blood has been reported in the literature for some time, yet the true origin of this is still the subject of much deliberation. The aim of this study was to investigate the phenomenon of a “blood microbiome” by providing a comprehensive description of bacterially-derived nucleic acids using a range of complementary molecular and classical microbiological techniques. For this purpose we utilised a set of plasma samples from healthy subjects (n = 5) and asthmatic subjects (n = 5). DNA-level analyses involved the amplification and sequencing of the 16S rRNA gene. RNA-level analyses were based upon the *de novo* assembly of unmapped mRNA reads and subsequent taxonomic identification. Molecular studies were complemented by viability data from classical aerobic and anaerobic microbial culture experiments. At the phylum level, the blood microbiome was predominated by Proteobacteria, Actinobacteria, Firmicutes and Bacteroidetes. The key phyla detected were consistent irrespective of molecular method (DNA vs RNA), and consistent with the results of other published studies. *In silico* comparison of our data with that of the Human Microbiome Project revealed that members of the blood microbiome were most likely to have originated from the oral or skin communities. To our surprise, aerobic and anaerobic cultures were positive in eight of out the ten donor samples investigated, and we reflect upon their source. Our data provide further evidence of a core blood microbiome, and provide insight into the potential source of the bacterial DNA / RNA detected in the blood. Further, data reveal the importance of robust experimental procedures, and identify areas for future consideration.

## Background

The term “microbiome” describes the genetic material encoding the various microbial populations that inhabit our body. In contrast, the term “microbiota” refers to the viable organisms that comprise these communities. Cell for cell, bacterial cells outnumber our own cells by a factor of ten, and the bacterial microbiome encodes for one hundred-fold more genes. The microbiome undertakes essential biological processes and thus it is unsurprising that a number of disease states are associated with changes in microbiome composition, termed “dysbiosis”. Whilst the colonisation of specific body sites in contact with the external environment (such as the gastrointestinal tract, skin and vagina) by microorganisms is both well-described and universally accepted ^1^, the existence of microbial populations in other “classically sterile” locations, including the blood, is a relatively new concept.

The presence of bacteria-specific DNA in the blood has been reported in the literature for some time, yet the true origin of this is still the subject of much deliberation. Mounting evidence supports the existence of a blood microbiome (*specifically, the presence of bacterial genetic material*) in humans ^2-11^ and various other species, including rodents, cats, chickens and cows ^12-15^. This has primarily been determined by amplification and sequencing of the bacterial 16S rRNA gene, or via whole genome sequencing. Such studies report the existence of bacteria-derived genetic material (DNA) within the circulation, but do not provide evidence for the presence of viable organisms.

Convention tells us that the blood is sterile in health, and bacteraemia, even at 1 – 10 bacterial cells per millilitre whole blood, is potentially life threatening. Despite this, several studies have presented evidence of bacteria or bacteria-like structures within the circulation in the absence of overt disease. It should be noted however that Martel and colleagues report that the bacteria-like particles often described following a range of imaging techniques represent non-living membrane vesicles and protein aggregates derived from the blood itself ^16^. McLaughlin surveyed the blood of 25 healthy donors and observed the presence of pleomorphic bacteria using dark-field microscopy, electron microscopy, polymerase chain reaction and fluorescent *in situ* hybridisation, in all samples analysed ^17^. Further, Potgieter and colleagues described the presence of blood-cell associated bacteria in a range of blood preparations using electron microscopic techniques ^8^. Significantly, Damgaard and colleagues found viable (*culturable*) bacteria in 62% of blood donations from donors with no overt disease ^18^. This finding is plausible given that various daily activities including chewing, tooth brushing and flossing result in the translocation of oral bacteria into the bloodstream ^19-22^, however one would expect such organisms to be rapidly targeted and removed with by the immune system in healthy individuals. Kell and colleagues provide a detailed hypothesis for the existence of the blood microbiome ^4,8^ and suggest that it is likely composed of organisms (or parts thereof) that translocate to the circulation from their usual place of habitation (classical niches such as the gastrointestinal tract, oral cavity, skin, vagina) – a process termed atopobiosis. They further describe how this could occur via well-described physiological processes including dendritic cells processes, via micro-fold cells, and in disease, via dysfunctional epithelial junctions. This explanation is supported by studies demonstrating a correlation between gut microbiota dysbiosis and altered microbial signatures detected in the blood ^23-25^, suggesting that the observed disease-associated blood microbiota is a consequence of increased bacterial translocation across the gut barrier. Furthermore, characterisation of the microbial populations in the coronary artery tissues by Lehtiniemi and colleagues ^26^ identified known members of the oral microbiota, suggesting that bacteria had translocated from the oral cavities into the bloodstream, potentially as a result of damage caused by tooth brushing or by leakage across the mucosal surfaces.

Various disease states are associated with blood microbiome dysbiosis ^2,6,24,25,27^, and these changes are likely reflective of dysbiosis at a distant site(s) with well-characterised microbial communities, and the result of subsequent translocation. Limited evidence also suggests that these changes may be disease-specific; Alzheimer’s disease for example, has been associated with the detection of mostly coccus microbes, whilst Parkinson’s disease has been associated with both coccus and bacillus bacteria ^8^. Such changes are of significant scientific and medical interest as they offer opportunities for biomarker and therapeutic target development ^25,27^.

Due to the long-held belief that the bloodstream of healthy individuals is sterile and since the blood is an unfavourable compartment for the microbes due to its bacteriostatic and bactericidal components ^1,13^, it is of principal significance to understand *whether* and *how* bacteria may persist in it. Accidental contamination during the collection of blood and or during downstream experimental procedures has been proposed as an alternative explanation for the existence of the blood microbiome *per se*. We support this explanation for the detection of *viable* bacteria within the bloodstream of healthy individuals, however, suggest that this phenomenon does not adequately explain the existence of the blood microbiome (*the presence of bacteria-derived nucleic acids*) when one considers the number of studies that demonstrate significant and apparently disease-specific differences in the composition of the blood microbiome. Moreover, examination of the bacterial taxa reported in these studies reveal similar blood microbiota compositions across the different studies, whereby Proteobacteria dominate (relative abundance values typically ranging from 85%-90%), and Firmicutes, Actinobacteria, and Bacteroidetes present to a lesser extent ^2,7,25,28^. This suggests the existence of a core blood microbiome profile that persists independent of study environment or analytical methodology.

Using a range of complementary molecular and classical molecular biology techniques, here we characterise the human blood microbiome in unparalleled detail; at the DNA level, we amplify and sequence the 16S rRNA gene, at the RNA level, we assemble almost 500,000,000 unmapped mRNA reads and map these to known taxa, and finally, we present viability data from classical aerobic and anaerobic microbial culture experiments. Our experimental approach is detailed in Figure 1.

**Figure 1.** Schematic representation of the multiple-method circulating microbiome characterisation approach implemented herein. NB: Biomarker and mechanistic data are not included within the scope of this publication and appear elsewhere.

## Materials and Methods

### Donor Samples

Atopic asthmatic individuals (n=5) with physician-diagnosed house dust mite allergy, and gender and age-matched healthy control subjects (n=5) were recruited to the study via SeraLabs Limited in accordance with the following criteria **(Table 1)**. Whole blood was drawn, following alcohol cleansing of the skin surface, into EDTA containing tubes and stored on ice prior to centrifugation at 1000×*g* to obtain the plasma component. All samples were analysed anonymously, and the authors obtained ethical approval (Keele University ERP3) and written informed consent to utilise the samples for the research reported herein.

**Table 1.**
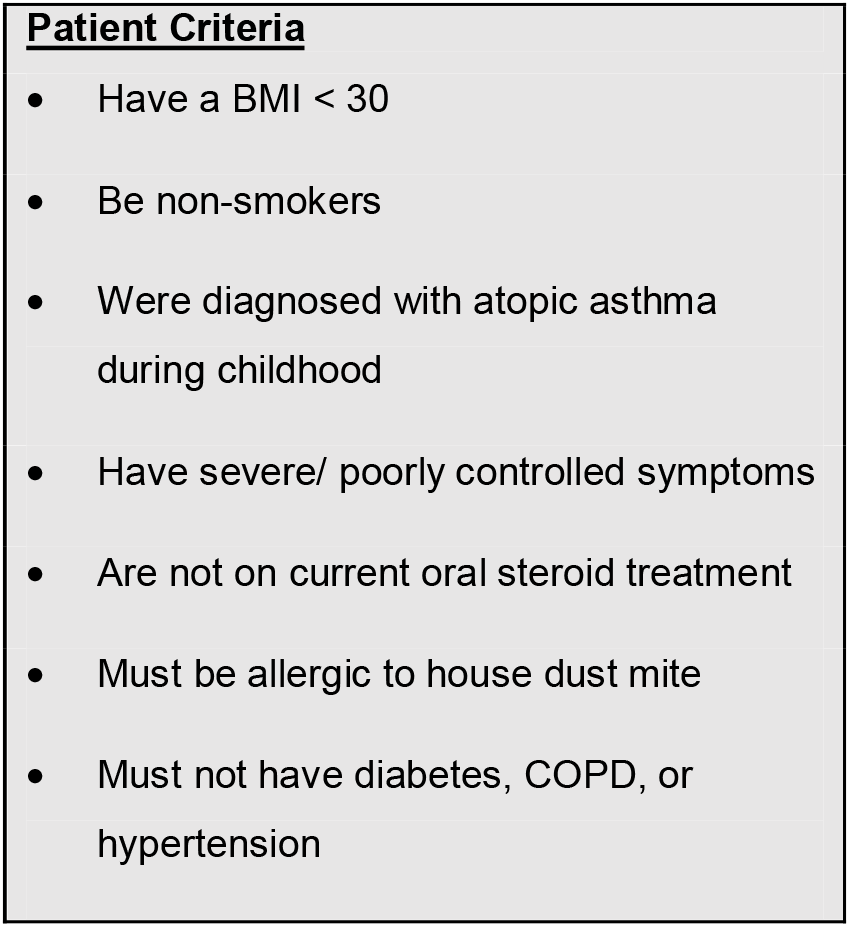
Donor population characteristics

### DNA-level: Meta-taxonomic Characterisation

The circulating microbiome was investigated at the DNA level by amplification and sequencing of the bacterial 16S ribosomal gene using oligonucleotide primers that target variable region 4 ^29^ **(Table 2)**. Direct amplification of the V4 region was performed using the Phusion Blood Direct kit (Thermofisher scientific) alongside a negative control reaction (in which blood was replaced with molecular biology grade water) that underwent the complete experimental procedure. Amplification was performed in triplicate as 20μl reactions containing; 1.0 μl plasma, 10μl 2X Phusion blood PCR buffer, 0.4μl Phusion Blood II DNA polymerase, 1.0 μl of each forward and reverse oligonucleotide primer (10μM), and 6.6μl of UV-treated molecular biology grade water. Cycling parameters were as follows; an initial 5 minutes denaturation step at 98°C followed by 33 cycles of; denaturation (1 second at 98°C), annealing (5 seconds at 55°C), and extension (15 seconds at 72°C), and a final extension at 72°C for 7 minutes.

Amplicons resulting from triplicate reactions were combined and purified using the MinElute PCR purification kit (Qiagen) prior to a further 7 cycles of PCR using AccuPrime *Pfx* SuperMix and a pair of V4 oligonucleotide primers we developed to incorporate the Illumina XT adapter in preparation for sequencing (**Table 2**). Cycling parameters were as follows; initial denaturation for 2 minutes at 95°C followed by 7 cycles of; denaturation (15 seconds at 95°C), annealing (30 seconds at 55°C), and extension (25 seconds at 68°C), and a final extension at 68°C for 10 minutes. PCR products were purified using the AMPure XP magnetic beads (Agencourt) at a ratio of 0.8 beads to sample (v/v), eluted in 20μl molecular biology grade water, and quantified using the Qubit 3.0 high-sensitivity DNA kit. Amplicons were barcoded using the Nextera DNA library kit, multiplexed, and sequenced using the Ilumina MiSeq system with a 250bp paired-end read metric. Bioinformatic analysis was performed using QIIME^30^ implemented as part of the Nephele 16S paired-end QIIME pipeline using closed reference clustering against the SILVA database ^31^ at a sequence identity of 99%. All other parameters remained as default.

**Table 2.**
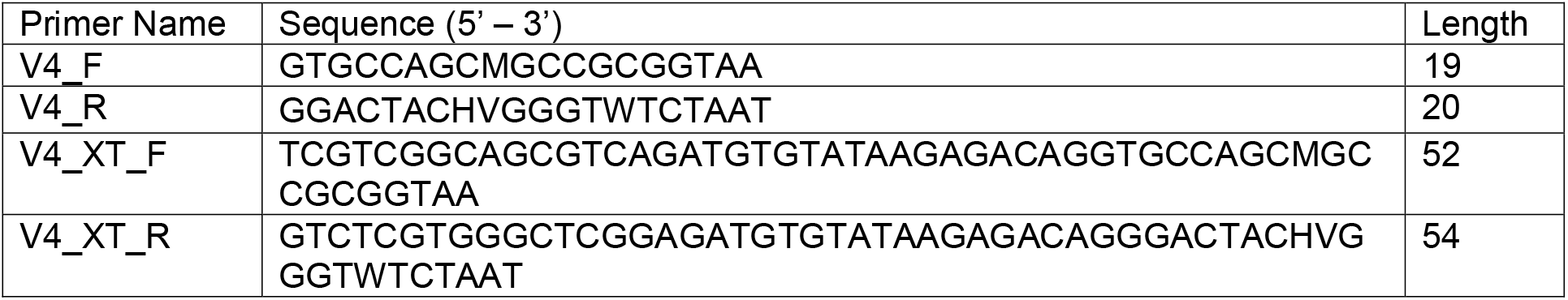
Oligonucleotide primer sequences

### RNA-level: De-novo Assembly of Unmapped RNA Reads

For the purposes of increasing our understanding of the molecular processes that are deregulated in patients with atopic asthma, we previously performed whole transcriptome sequencing on RNA extracted from each donor plasma sample (data currently unpublished). Here, we hypothesised that the non-mapping (likely non-human) reads that often result from such analyses would represent microbial community members that were present in the blood at the time of RNA extraction. To this end, RNA reads were mapped to the *Homo sapiens* genome version hg38 using Tophat with default parameters ^32^. Reads failing to map to hg38 were identified from the resulting BAM file and reads, in fastq format, extracted using bedtools bamtofastq. In order to streamline our strategy, a single de novo transcriptome was produced by concatenating all of the unmapped reads produced from the entire study, and assembling these using Trinity ^33^ to form features representative of candidate non-human genes. To increase computational efficiency, the total read population was subsampled to a depth of 1M, 10M, 25M, and 100M, and the number of reads that these features explained computed by building a bowtie2 index out of each resulting transcriptome, and mapping the unmapped read population from each sample back to it. The transcriptome assembly with the highest mapping rate was used for further analysis as follows; (1) Abundance estimation – the transcriptome was indexed for bowtie2 and the number of reads mapping to each feature determined for each of our donor samples using RSEM ^34^. (2) A matrix of expression values was produced using the abundance_estimates_to_matrix.pl script packaged with Trinity ^35^. (3) Statistical analysis - a differential expression analysis was conducted to identify candidate non-human genes that were significantly differentially expressed between the disease and disease-free donors using edgeR ^36^. (4) Identification - the taxonomic identity of each assembled feature was determined using Kraken ^37^ and visualised using Pavian ^38^.

### Classical Culture: Bacterial Viability

Classical microbiological culture, using a range of substrates, was carried out to determine whether the human plasma samples contained any viable bacterial cells, i.e. those capable of proliferation. For each sample, 250μl plasma was inoculated into 9ml of brain heart infusion broth and incubated for 5 days at 37°C. For each culture a negative control was generated, whereby 9ml of brain heart infusion broth was inoculated with 250μl of ultra-violet sterilised distilled water and incubated for 5 days at 37°C. The inoculated broth was plated onto agar plates (Columbia agar + 5% horse blood; CLED medium; A.R.I.A + horse blood) and incubated under either aerobic (Columbia blood agar; CLED medium) or anaerobic (A.R.I.A agar) conditions at 37°C for a minimum of three days. Bacterial growth was evaluated by sight, and a selection of individual colonies from each plate were selected for identification by total 16S gene amplification and Sanger sequencing.

## Results and Discussion

### Donor Characteristics

The donor population were all female and all “never smokers”. The asthmatic population were 39.6 years in age (range 19 – 52) with a mean body mass index of 24.4 (range 21.5 – 27.8) and all had physician-diagnosed atopic asthma resulting from house dust mite allergy. The control population were 39.4 years in age (range 23 – 49) with a mean body mass index of 24.3 (range 21.0 – 26.4) and were disease-free. There were no statistically significant differences in age (P = 0.98) or BMI (P = 0.93) between the two groups **(Table 3)**.

**Table 3.** Donor Population Characteristics

| Donor | Age (years) | BMI (kg/m<sup>2</sup>) | Smoking History | Diagnosis |
| --- | --- | --- | --- | --- |
| BRH1017873 | 50-60 | 27.8 | Never | Atopic Asthma, HDM |
| BRH1017874 | 30-40 | 27.3 | Never | Atopic Asthma, HDM |
| BRH1017875 | 40-50 | 23.3 | Never | Atopic Asthma, HDM |
| BRH1017876 | 10-20 | 21.5 | Never | Atopic Asthma, HDM |
| BRH1017877 | 40-50 | 22.3 | Never | Atopic Asthma, HDM |
| BRH1017878 | 40-50 | 26.4 | Never | Healthy |
| BRH1017879 | 20-30 | 21 | Never | Healthy |
| BRH1017880 | 40-50 | 26.4 | Never | Healthy |
| BRH1017881 | 40-50 | 24.9 | Never | Healthy |
| BRH1017882 | 30-40 | 22.7 | Never | Healthy |

### DNA-level Circulating Microbiome – Community Structure

The presence of bacterial DNA within the blood of our study cohort was evaluated by amplification and sequencing of the 16S RNA gene variable region 4. A negative experimental control sample mirrored our study samples through the entire experimental procedure downstream of venepuncture. This involved using ultra-violet treated distilled water in replacement of human blood during PCR amplification of the 16S rRNA V4 region. The negative control PCR product was then submitted to all downstream applications that the human blood underwent. This included bead-based purification of the 16S rRNA V4 amplicons, agarose gel electrophoresis, XT-tagging, library preparation and sequencing of the 16S rRNA V4 amplicons. Despite this control reaction being PCR-negative, it was nevertheless submitted for sequencing.

Using QIIME^30^ implemented as part of Nephele, a total of 243, 853 sequencing reads from the amplified V4 region passed quality assessment (mean = 24,385 reads per sample; range = 10, 742 – 35, 701 reads). The results of our taxonomic classification are shown in **(Figure 2 A-D)** at the phylum and genus levels.

**Figure 2.** Relative abundance of the most abundant taxa (>1%) as determined by amplification and sequencing of the 16S rRNA gene variable region 4. Data are mean abundance expressed as a percentage of the total bacterial sequence count. (A) Phylum-level data grouped by condition, (B) Genus-level data grouped by condition, (C) Phylum-level individual sample data, (D) Genus-level individual sample data.

Our negative control reaction generated a small number of reads that were identified as the following genera; *Halomonas (6 reads)*, *Corynebacterium (64)*, *Staphylococcus (24)*, *Ralstonia (1726)*, *Stenotrophomonas (460)*, *Pseudomonas (276)*, *Escherichia*-*Shigella (2420)* and *Ruminococcus* (405), but was overwhelmingly composed of reads mapping to the genus *Serratia (18000)*. These genera have been reported previously as contaminants of next generation sequencing experiments ^39,40^ but importantly here, were either distinct from the taxa identified within our samples, or present at much lower levels.

At the phylum level, the blood microbiome was predominated by Proteobacteria (88% of all bacterial DNA in the control population, and 80.9% in the asthmatic population) followed by Actinobacteria (control = 7.8%, asthmatic = 7.1%), Firmicutes (control = 3.5%, asthmatic = 9.2%) and Bacteroidetes (control = 0.1%, asthmatic = 2.2%). These findings mirror previous studies ^2,7,25,28^ and further support the notion of a core blood microbiome predominated by four key phyla.

At the genus level, our blood samples were predominated by the genus *Achromobacter ^41^* which accounted for 51.1% and 45.3% of the total bacterial DNA detected in the control and asthma donors respectively. To a lesser extent, the blood samples also comprised members of the *Pseudomonas* (12.8%, 7.5%) ^7,9,18^, *Serratia\** (0.9%, 11.6%) ^42^, *Sphingomonas* (3.8%, 5.1%) ^7,25^, *Staphylococcus* (5.5%, 2.8%) ^7,18,43,44^, *Corynebacterium* (3.2%, 5.5%) ^7,9^, and *Acinetobacter* (3.7%, 2.8%) ^7^ genera. The genus *Serratia* was excluded from further analysis as the study samples presented with less reads than did the corresponding negative control reaction and thus it was considered a contaminant.

Whilst all genera found have been previously described in the blood (see references), the predominance of *Achromobacter,* which is not classically associated with the blood microbiome, warrants further consideration. Indeed, *Achromobacter* has been detected abundantly in the lower respiratory tract of healthy mice ^45^, humans (HPM airway dataset), and in various respiratory conditions ^46,47^. Furthermore, no *Achromobacter* was detected in our experimental control reactions suggesting that its presence is not the result of experimental contamination.

Although differential composition of the blood microbiome in response to pathology was not the main focus of this study, we performed principal coordinates analysis and despite the relatively small sample set, this revealed good separation between the two treatment groups based upon their microbiome profile, suggesting that the blood microbiome community was altered in the asthmatic subjects (**Figure 3**).

**Figure 3.** Principal coordinates analysis of weighted unifrac distances for control (blue) and asthmatic (red) blood microbiome profiles. Each dot represents an individual sample, and the microbiome of samples that appear more closely together are more similar.

### DNA-level Circulating Microbiome – Likely Origins

In accordance with our hypothesis that the blood microbiome exists as a consequence of bacterial translocation from other microbiome niches within the body, we compared the data generated herein with gastrointestinal tract, oral cavity, and skin data made available by the Human Microbiome Project (HMP). In each case, the HMP data and our own were combined, weighted UniFrac distances calculated, and a principal coordinates analysis performed. **Figure 4** demonstrates that the blood microbiome of our control and asthmatic donors clustered more closely in PCoA space with the oral cavity and skin HMP data, than it did with the gastrointestinal tract HMP data. This suggests that the blood microbiome community is perhaps more likely to result from the translocation of organisms from the oral cavity and skin, than from organisms that colonise the gastrointestinal tract.

**Figure 4.** Principal coordinates analysis of weighted unifrac distances between variable region 4 16S sequencing data from our donors and the Human Microbiome Project (A) Gut, (B) Oral Cavity and (C) Skin data. Each dot represents an individual sample, and the microbiome of samples that appear more closely together are more similar. In each case, our control donor samples appear in blue, and our asthmatic donor samples appear in red. Further sample details are provided beneath each figure, and the number of datasets representing each anatomic location is provided in brackets.

Various daily activities including chewing, tooth brushing and flossing have been shown to result in the translocation of bacteria from the oral cavity into the bloodstream ^19-22^. Further, the skin has a distinct microbial community and is susceptible to injury, and thus represents a large surface area through which such organisms may translocate to the bloodstream. It is important here to consider sources of contamination; venepuncture, the process through which the majority of blood samples are obtained, is recognised as a cause of transient bacteraemia ^48^, and despite the use of preventative measures such as alcohol cleansing of the skin prior to breaking the surface, there remains the possibility that organisms entered the sample from the skin via this route.

One important, but often overlooked limitation of DNA-based microbiome characterisation is that DNA persists post-mortem, even in the presence of harsh environmental conditions. From such analyses it is therefore impossible to confirm whether an organism is present and viable, is present but non-viable, or whether the organism has since left the environment in question yet it’s DNA persists.

### RNA-level Circulating Microbiome – Community Structure

We hypothesise that some of the non-mapping (likely non-human) reads that often result from whole transcriptome analyses (RNA-seq) represent microbial community members that were present (or parts thereof) in the blood at the time of RNA extraction. Furthermore, given the unstable nature of extracellular circulatory RNA^49^ in addition to the presence of circulatory ribonucleases that actively degrade RNA ^47^,, we suggest that the detection of bacterial RNA goes further towards confirming the recent presence of these microbes within the blood when compared with DNA-based approaches. From our previous studies of the circulating transcriptome of our donor community, a total of 439,448,931 paired RNA reads failed to map to the human genome. These reads were used for the following analyses as randomly subsampled populations of 1 million, 10 million, 25 million and 100 million read pairs. Mapping the total read population back to the subsampled populations allowed us to assess how well each subsampled population approximated the starting (entire, ~ 440M read) population. Data revealed only marginal improvements in whole community representation as the subsampled population increased (1M, 10M, 25M, 100M represented 65.05, 66.05, 66.81, 64.24% of the total read population). A subsampled population of 25 million reads (25M) provided an acceptable balance between read representation and computational efficiency and was therefore used for transcriptome assembly. The transcriptome comprised 2050 candidate “non-human genes” with a mean GC content of 53%. Ten-percent of these genes were greater than 517bp, and over half were at least 263bp. Taxonomic identification of each assembled feature was determined using Kraken ^37^ and this revealed that 729 of the 2050 features were of bacterial origin and 7 features were from archaea (pertaining to the taxa Thermoplasmata, which has been previously associated with the human microbiome ^50^) (**Figure 5**). Although we identified 13 features of apparent viral origin, we did not consider these in any further detail given they appeared to pertain to the Moloney murine leukemia virus, a commonly utilised reverse transcriptase used in molecular procedures. It should be noted that the Kraken database does not include fungi, and therefore this kingdom was not represented within our data.

**Figure 5.** Taxonomic classification of each feature assembled from unmapped RNA sequencing reads using Trinity and identified using Kraken. The numbers present by each taxonomic classification refer to the number of features classified as such (e.g. 379 assembled features were identified as Proteobacteria). D – domain, P – phylum, F – family, G – genus, S – species.

At the phylum level, the whole transcriptome data was predominated by assembled Proteobacteria sequences (379 sequences, 52%), followed by Firmicutes (143, 19.8%), Actinobacteria (112, 15.5%) and Bacteroidetes (35, 4.8%). In considering the total number of reads mapping to each feature, 379.4M reads mapped to bacterial features (out of a total of 395M reads). Of those reads mapping to bacterial-derived sequences, Proteobacteria (74.9%, 47.0%; Control, Asthma) and Firmicutes (19.5%, 48.0%) predominated with, Actinobacteria (0.01%, 0.04%) and Bacteroidetes (0.05%, 0.008%) present to a much lesser extent. These findings support our DNA-level phylum data and mirror previous studies ^2,7,25,28^.

At the genus level, the whole transcriptome data was predominated by the genera *Paenibacillus* (17.8%, 44.6%; Control, Asthma reads)^51^, *Escherichia* (11.8%, 13.1%) 7,9,43,52, *Acinetobacter* (0.7%, 0.4%) ^7^, *Pseudomonas* (0.8%, 0.1%) ^7,9,18,53^ and *Propionibacterium* (0.5%, 0.2%) ^9,18,44,53^. With the exception of *Paenibacillus,* all genera detected via *de novo* transcriptome assembly were also present in our DNA-level data and all genera found have been previously described in the blood (see references). The fact that *Paenibacillus* was present within our RNA-level analyses yet absent from our DNA-level data led us to consider whether it could have been introduced as a contaminant during the RNA extraction, library preparation or sequencing procedures. Given that negative control reactions are not routinely conducted for RNA-seq type applications, we were unable to experimentally confirm this. However, we did identify from the literature that this genus has been reported as a common reagent and laboratory contaminant, albeit at the DNA level ^40^. Nevertheless there appeared to be little consistency in its presence within our sample set (mean % of reads 39.5 ± 40.3%) and we noted a difference in abundance between our experimental groups despite all preparation procedures being the same. The exact source of this RNA thus remains open to speculation.

### Classical Culture – Presence of Viable Organisms

The presence of viable, proliferating bacteria in the blood was assessed using growth culture assays as previously described. Bacterial cultures were positive for 80% of blood samples assayed (8 out of 10 samples; 4 control blood samples and 4 asthma blood samples), whilst all negative control plates had no growth as expected. These results are relatively consistent with previous studies, whereby 2-100% of blood samples were positive for bacterial growth ^18, 54-57^. Unexpectedly, bacterial growth was observed in aerobic conditions for all culture-positive blood samples, but anaerobic growth was only observed for four of the culture-positive blood samples. This is contradictory to previous studies, where bacterial growth from blood-cultures has predominately been achieved using anaerobic conditions ^18,55^.

In all instances, bacterial growth was monoculture on microscopy and thus 16S colony PCR (amplifying the entire 16S rRNA gene) and Sanger sequencing was conducted on a three independent colonies per plate for identification purposes. Bacteria were identified using Sanger sequencing followed by classification with Kraken. Bacteria isolated from the aerobic cultures included the following genera; *Staphylococcus* (49 sequences), *Micrococcus* (12), *Kocuria* (6), *Corynebacterium* (6) and *Propionibacterium* (1). Bacteria isolated from the anaerobic cultures were less variable and included members from the facultatively anaerobic *Staphylococcus* genus only (**Figure 6**). These genera belong to the phyla Actinobacteria (*Corynebacterium*, *Kocuria*, *Micrococcus*) and Firmicutes (*Staphylococcus*) and were all represented in our 16S DNA level data (**Figure 4**) and detected within our RNA data. Individual sample data is presented in (**Table 4**). It is noteworthy that, with the exception of *Kocuria*, all bacteria identified displayed some of the highest total relative abundance scores in the 16S sequencing results; *Corynebacterium* (4.2%), *Kocuria* (0.2%), *Micrococcus* (1.30%), and *Staphylococcus* at 4.3%. Due to the long-held belief that the bloodstream of healthy individuals is sterile and since the blood is an unfavourable compartment for the microbes due to its bacteriostatic and bactericidal components; here we consider the likely source of these viable organisms. The skin microbiome is dominated by members of the genera *Corynebacterium*, *Micrococcus*, *Staphylococcus*, and *Propionibacterium*, the proportions of which vary markedly between individuals ^58^. Furthermore, several studies report the presence of the genus *Kocuria* on the skin of humans and other mammals ^59-61^. We therefore suggest that the organisms detected through our microbial culture experiments most likely originate from the skin. Whilst transient bacteraemia due to a breach of the skin barrier is an accepted occurrence, one would expect such organisms to be rapidly targeted and removed by the immune system.

**Figure 6.** Taxonomic classification of total 16S data generated by colony PCR and Sanger sequencing. The numbers present by each taxonomic classification indicate the number of colonies that were identified with that identity. D – domain, P – phylum, F – family, G – genus, S – species.

**Table 4.** Identification of cultured organisms using 16S colony PCR and Sanger Sequencing. Culture Neg denotes no bacterial growth on the substrates used.

| Growth<br>Conditio<br>n | Control Samples |  |  |  |  | Asthma Samples |  |  |  |
| --- | --- | --- | --- | --- | --- | --- | --- | --- | --- |
|  | BRH101787<br>8 | BRH101787<br>9 | BRH1017880 | BRH1017881 | BRH101788<br>2 | BRH1017874 | BRH101787<br>5 | BRH1017876 | BRH1017877 |
| Aerobic<br>Growth | *Kocuria*<br>*rhizophila* | *Micrococcu*<br>*s luteus* | *Staphylococcus*<br>*haemolyticus*<br>*Staphylococcus*<br>*epidermidis*<br>*Propionibacteriu*<br>*m acnes* | *Staphylococc*<br>*us*<br>*haemolyticus* | Culture Neg | *Staphylococc*<br>*us*<br>*haemolyticus* | Culture Neg | *Corynebacteriu*<br>*m halotolerans* | *Staphylococc*<br>*us epidermidis* |
| Anaerobi<br>c Growth | Culture Neg | Culture Neg | *Staphylococcus*<br>*epidermidis* | *Staphylococc*<br>*us hominis* | Culture Neg | Culture Neg | Culture Neg | Culture Neg | *Staphylococc*<br>*us epidermidis* |

We therefore suggest that the viable organisms detected through classical microbial culture analysis are the result of venepuncture contamination whereby organisms from the skin are drawn into the vacutainer, contaminating the sample. An alternative hypothesis suggests that these bacteria were present in the blood in a dormant state (i.e. not contaminants), and were somehow revived following pre-growth in brain heart infusion broth prior to plating (see Kell et al for a detailed description of this hypothesis ^8^), however, this hypothesis is still under intense investigation.

### Concluding Remarks

This study utilised a range of molecular and classical microbiology approaches to characterise the human blood microbiome in unparalleled detail. Our DNA and RNA-based studies revealed a diverse community of bacteria, the main members of which having been described in a range of other studies and therefore providing further evidence of a core blood microbiome. Although disease associated changes in the blood microbiome were not the focus of this study, the fact we identified such changes is encouraging and supports efforts to identify circulating microbiome signatures indicative of disease.

Whilst we attribute the finding of viable organisms in our plasma samples to venepuncture-associated contamination of the blood sample (and make recommendations to avoid this in future) and or the phenomena of dormancy, the presence of these viable organisms does not undermine our exciting molecular data that reports an abundance of bacteria-associated DNA and RNA within the blood, likely present due to translocation from classical microbiome niches (such as the gut, oral cavity and skin), and with the clear potential to be developed as a biomarker of microbiome status at distant anatomical sites.

On reflecting upon our experimental approach, we make the following recommendations for future studies;

(1) Significant attention should be paid to blood collection procedures as any contamination occurring at this stage impacts upon all downstream procedures. In addition to alcohol cleansing of the skin (as performed in this study), we recommend that the first volume drawn is diverted to a secondary tube, and analysed separately. This will allow investigation of the contribution that venepuncture-associated contamination makes, and allow robust analysis of the dormancy hypothesis.

(2) The inclusion of negative control reactions that are subject to the whole range of experimental procedures, including library preparation and sequencing, is absolutely essential.

(3) Where possible, RNA-seq studies used for unmapped read assembly should include negative control samples that are subject to all of the experimental procedures alongside the study samples. This will allow an appraisal of how significant reagent / laboratory procedure contamination is in these studies. Often the use of RNA-seq derived unmapped reads for microbiome characterisation is a “secondary use”, and this is simply not possible.

(4) Seminal studies are still required to satisfactorily investigate the phenomenon of bacterial translocation from well-characterised microbiome niches to the blood.

## Acknowledgements

This study used the Nephele platform from the National Institute of Allergy and Infectious Diseases (NIAID) Office of Cyber Infrastructure and Computational Biology (OCICB) in Bethesda, MD. This article was published as a pre-print with the following DOI: https://doi.org/10.1101/359760 ^62^

## Data Availability

The sequencing data utilised in this project can be found in the Sequence Read Archive, NIH, under the identifier SUB4654957.

## References

1. Markova, N. D. L-form bacteria cohabitants in human blood: significance for health and diseases. Discov Med 23, 305-313 (2017).

2. Amar, J. et al. Blood microbiota dysbiosis is associated with the onset of cardiovascular events in a large general population: the D.E.S.I.R. study. PLoS One 8, e54461, doi:10.1371/journal.pone.0054461 (2013).

3. Bhattacharyya, M., Ghosh, T., Shankar, S. & Tomar, N. The conserved phylogeny of blood microbiome. Mol Phylogenet Evol 109, 404-408, doi:10.1016/j.ympev.2017.02.001 (2017).

4. Kell, D. B. & Pretorius, E. On the translocation of bacteria and their lipopolysaccharides between blood and peripheral locations in chronic, inflammatory diseases: the central roles of LPS and LPS-induced cell death. Integr Biol (Camb) 7, 1339-1377, doi:10.1039/c5ib00158g (2015).

5. Li, Q. et al. Identification and Characterization of Blood and Neutrophil-Associated Microbiomes in Patients with Severe Acute Pancreatitis Using Next-Generation Sequencing. Front Cell Infect Microbiol 8, 5, doi:10.3389/fcimb.2018.00005 (2018).

6. Mangul, S. et al. Total RNA Sequencing reveals microbial communities in human blood and disease specific effects. bioRxiv, doi:10.1101/057570 (2016).

7. Païssé, S. et al. Comprehensive description of blood microbiome from healthy donors assessed by 16S targeted metagenomic sequencing. Transfusion 56, 1138-1147, doi:10.1111/trf.13477 (2016).

8. Potgieter, M., Bester, J., Kell, D. B. & Pretorius, E. The dormant blood microbiome in chronic, inflammatory diseases. FEMS Microbiol Rev 39, 567-591, doi:10.1093/femsre/fuv013 (2015).

9. Dinakaran, V. et al. Elevated levels of circulating DNA in cardiovascular disease patients: metagenomic profiling of microbiome in the circulation. PLoS One 9, e105221, doi:10.1371/journal.pone.0105221 (2014).

10. Rajendhran, J., Shankar, M., Dinakaran, V., Rathinavel, A. & Gunasekaran, P. Contrasting circulating microbiome in cardiovascular disease patients and healthy individuals. Int J Cardiol 168, 5118-5120, doi:10.1016/j.ijcard.2013.07.232 (2013).

11. Nikkari, S., McLaughlin, I. J., Bi, W., Dodge, D. E. & Relman, D. A. Does blood of healthy subjects contain bacterial ribosomal DNA? J Clin Microbiol 39, 1956-1959, doi:10.1128/JCM.39.5.1956-1959.2001 (2001).

12. Vientós-Plotts, A. I. et al. Dynamic changes of the respiratory microbiota and its relationship to fecal and blood microbiota in healthy young cats. PLoS One 12, e0173818, doi:10.1371/journal.pone.0173818 (2017).

13. Mandal, R. K. et al. An investigation into blood microbiota and its potential association with Bacterial Chondronecrosis with Osteomyelitis (BCO) in Broilers. Sci Rep 6, 25882, doi:10.1038/srep25882 (2016).

14. Sze, M. A. et al. Changes in the bacterial microbiota in gut, blood, and lungs following acute LPS instillation into mice lungs. PLoS One 9, e111228, doi:10.1371/journal.pone.0111228 (2014).

15. Jeon, S. J. et al. Blood as a route of transmission of uterine pathogens from the gut to the uterus in cows. Microbiome 5, 109, doi:10.1186/s40168-017-0328-9 (2017).

16. Martel, J., Wu, C. Y., Huang, P. R., Cheng, W. Y. & Young, J. D. Pleomorphic bacteria-like structures in human blood represent non-living membrane vesicles and protein particles. Sci Rep 7, 10650, doi:10.1038/s41598-017-10479-8 (2017).

17. McLaughlin, R. W. et al. Are there naturally occurring pleomorphic bacteria in the blood of healthy humans? J Clin Microbiol 40, 4771-4775 (2002).

18. Damgaard, C. et al. Viable bacteria associated with red blood cells and plasma in freshly drawn blood donations. PLoS One 10, e0120826, doi:10.1371/journal.pone.0120826 (2015).

19. Forner, L., Larsen, T., Kilian, M. & Holmstrup, P. Incidence of bacteremia after chewing, tooth brushing and scaling in individuals with periodontal inflammation. J Clin Periodontol 33, 401-407, doi:10.1111/j.1600-051X.2006.00924.x (2006).

20. Horliana, A. C. et al. Dissemination of periodontal pathogens in the bloodstream after periodontal procedures: a systematic review. PLoS One 9, e98271, doi:10.1371/journal.pone.0098271 (2014).

21. Parahitiyawa, N. B., Jin, L. J., Leung, W. K., Yam, W. C. & Samaranayake, L. P. Microbiology of odontogenic bacteremia: beyond endocarditis. Clin Microbiol Rev 22, 46-64, Table of Contents, doi:10.1128/CMR.00028-08 (2009).

22. Lockhart, P. B. et al. Bacteremia associated with toothbrushing and dental extraction. Circulation 117, 3118-3125, doi:10.1161/CIRCULATIONAHA.107.758524 (2008).

23. Ono, S. et al. Detection of microbial DNA in the blood of surgical patients for diagnosing bacterial translocation. World J Surg 29, 535-539, doi:10.1007/s00268-004-7618-7 (2005).

24. Sato, J. et al. Gut dysbiosis and detection of †live gut bacteria† in blood of Japanese patients with type 2 diabetes. Diabetes Care 37, 2343-2350, doi:10.2337/dc13-2817 (2014).

25. Lelouvier, B. et al. Changes in blood microbiota profiles associated with liver fibrosis in obese patients: A pilot analysis. Hepatology 64, 2015-2027, doi:10.1002/hep.28829 (2016).

26. Lehtiniemi, J., Karhunen, P. J., Goebeler, S., Nikkari, S. & Nikkari, S. T. Identification of different bacterial DNAs in human coronary arteries. Eur J Clin Invest 35, 13-16, doi:10.1111/j.1365-2362.2005.01440.x (2005).

27. Ling, Z. et al. Blood microbiota as a potential noninvasive diagnostic biomarker for liver fibrosis in severely obese patients: Choose carefully. Hepatology 65, 1775-1776, doi:10.1002/hep.28987 (2017).

28. Olde Loohuis, L. M. et al. Transcriptome analysis in whole blood reveals increased microbial diversity in schizophrenia. Transl Psychiatry 8, 96, doi:10.1038/s41398-018-0107-9 (2018).

29. Kozich, J. J., Westcott, S. L., Baxter, N. T., Highlander, S. K. & Schloss, P. D. Development of a dual-index sequencing strategy and curation pipeline for analyzing amplicon sequence data on the MiSeq Illumina sequencing platform. Appl Environ Microbiol 79, 5112-5120, doi:10.1128/AEM.01043-13 (2013).

30. Caporaso, J. G. et al. QIIME allows analysis of high-throughput community sequencing data. Nat Methods 7, 335-336, doi:10.1038/nmeth.f.303 (2010).

31. Quast, C. et al. The SILVA ribosomal RNA gene database project: improved data processing and web-based tools. Nucleic Acids Res 41, D590-596, doi:10.1093/nar/gks1219 (2013).

32. Trapnell, C. et al. Differential gene and transcript expression analysis of RNA-seq experiments with TopHat and Cufflinks. Nat Protoc 7, 562-578, doi:10.1038/nprot.2012.016 (2012).

33. Haas, B. J. et al. De novo transcript sequence reconstruction from RNA-seq using the Trinity platform for reference generation and analysis. Nat Protoc 8, 1494-1512, doi:10.1038/nprot.2013.084 (2013).

34. Li, B. & Dewey, C. N. RSEM: accurate transcript quantification from RNA-Seq data with or without a reference genome. BMC Bioinformatics 12, 323, doi:10.1186/1471-2105-12-323 (2011).

35. Grabherr, M. G. et al. Full-length transcriptome assembly from RNA-Seq data without a reference genome. Nat Biotechnol 29, 644-652, doi:10.1038/nbt.1883 (2011).

36. Robinson, M. D., McCarthy, D. J. & Smyth, G. K. edgeR: a Bioconductor package for differential expression analysis of digital gene expression data. Bioinformatics 26, 139-140, doi:10.1093/bioinformatics/btp616 (2010).

37. Wood, D. E. & Salzberg, S. L. Kraken: ultrafast metagenomic sequence classification using exact alignments. Genome Biol 15, R46, doi:10.1186/gb-2014-15-3-r46 (2014).

38. Breitwieser, F. P. & Salzberg, S. L. Pavian: Interactive analysis of metagenomics data for microbiomics and pathogen identification. bioRxiv, doi:10.1101/084715 (2016).

39. Laurence, M., Hatzis, C. & Brash, D. E. Common contaminants in next-generation sequencing that hinder discovery of low-abundance microbes. PLoS One 9, e97876, doi:10.1371/journal.pone.0097876 (2014).

40. Salter, S. J. et al. Reagent and laboratory contamination can critically impact sequence-based microbiome analyses. BMC Biol 12, 87, doi:10.1186/s12915-014-0087-z (2014).

41. Gyarmati, P. et al. Metagenomic analysis of bloodstream infections in patients with acute leukemia and therapy-induced neutropenia. Sci Rep 6, 23532, doi:10.1038/srep23532 (2016).

42. Moriyama, K. et al. Polymerase chain reaction detection of bacterial 16S rRNA gene in human blood. Microbiol Immunol 52, 375-382, doi:10.1111/j.1348-0421.2008.00048.x (2008).

43. Such, J. et al. Detection and identification of bacterial DNA in patients with cirrhosis and culture-negative, nonneutrocytic ascites. Hepatology 36, 135-141, doi:10.1053/jhep.2002.33715 (2002).

44. Marques da Silva, R., Lingaas, P. S., Geiran, O., Tronstad, L. & Olsen, I. Multiple bacteria in aortic aneurysms. J Vasc Surg 38, 1384-1389, doi:10.1016/S0741 (2003).

45. Singh, N., Vats, A., Sharma, A., Arora, A. & Kumar, A. The development of lower respiratory tract microbiome in mice. Microbiome 5, 61, doi:10.1186/s40168-017-0277-3 (2017).

46. Hogan, D. A. et al. Analysis of Lung Microbiota in Bronchoalveolar Lavage, Protected Brush and Sputum Samples from Subjects with Mild-To-Moderate Cystic Fibrosis Lung Disease. PLoS One 11, e0149998, doi:10.1371/journal.pone.0149998 (2016).

47. Segal, L. N., Rom, W. N. & Weiden, M. D. Lung microbiome for clinicians. New discoveries about bugs in healthy and diseased lungs. Ann Am Thorac Soc 11, 108-116, doi:10.1513/AnnalsATS.201310-339FR (2014).

48. Depcik-Smith, N. D., Hay, S. N. & Brecher, M. E. Bacterial contamination of blood products: factors, options, and insights. J Clin Apher 16, 192-201 (2001).

49. Tsui, N. B., Ng, E. K. & Lo, Y. M. Stability of endogenous and added RNA in blood specimens, serum, and plasma. Clin Chem 48, 1647-1653 (2002).

50. Lurie-Weinberger, M. N. & Gophna, U. Archaea in and on the Human Body: Health Implications and Future Directions. PLoS Pathog 11, e1004833, doi:10.1371/journal.ppat.1004833 (2015).

51. Sáez-Nieto, J. A. et al. spp. isolated from human and environmental samples in Spain: detection of 11 new species. New Microbes New Infect 19, 19-27, doi:10.1016/j.nmni.2017.05.006 (2017).

52. Francés, R. et al. A sequential study of serum bacterial DNA in patients with advanced cirrhosis and ascites. Hepatology 39, 484-491, doi:10.1002/hep.20055 (2004).

53. Bhatt, A. S. et al. In search of a candidate pathogen for giant cell arteritis: sequencing-based characterization of the giant cell arteritis microbiome. Arthritis Rheumatol 66, 1939-1944, doi:10.1002/art.38631 (2014).

54. Domingue, G. J. & Schlegel, J. U. Novel bacterial structures in human blood: cultural isolation. Infect Immun 15, 621-627 (1977).

55. Jiménez, E. et al. Isolation of commensal bacteria from umbilical cord blood of healthy neonates born by cesarean section. Curr Microbiol 51, 270-274, doi:10.1007/s00284-005-0020-3 (2005).

56. Wilson, W. R., Van Scoy, R. E. & Washington, J. A. Incidence of Bacteremia in Adults Without Infection. Journal of Clinical Microbiology 2, 94-95 (1975).

57. Panaiotov, S. Cultural Isolation and Characteristics of the Blood Microbiome of Healthy Individuals. Scientific Research 008, 406-421 (2018).

58. Tett, A. et al. Unexplored diversity and strain-level structure of the skin microbiome associated with psoriasis. NPJ Biofilms Microbiomes 3, 14, doi:10.1038/s41522-017-0022-5 (2017).

59. Cosseau, C. et al. Proteobacteria from the human skin microbiota: Species-level diversity and hypotheses. One Health 2, 33-41, doi:10.1016/j.onehlt.2016.02.002 (2016).

60. McIntyre, M. K., Peacock, T. J., Akers, K. S. & Burmeister, D. M. Initial Characterization of the Pig Skin Bacteriome and Its Effect on In Vitro Models of Wound Healing. PLoS One 11, e0166176, doi:10.1371/journal.pone.0166176 (2016).

61. Grice, E. A. et al. A diversity profile of the human skin microbiota. Genome Res 18, 1043-1050, doi:10.1101/gr.075549.107 (2008).

62. Whittle, E., Leonard, M., Harrison, R., Gant, T. & Tonge, D. Multi-Method Characterisation of the Human Circulating Microbiome. BioRxiv, doi:https://doi.org/10.1101/359760 (2018).

